# Comprehensive Modeling of Spinal Muscular Atrophy in *Drosophila melanogaster*

**DOI:** 10.1101/394908

**Authors:** Ashlyn M. Spring, Amanda C. Raimer, Christine D. Hamilton, Michela J. Schillinger, A. Gregory Matera

## Abstract

Spinal muscular atrophy (SMA) is a neurodegenerative disorder that affects motor neurons, primarily in young children. SMA is caused by mutations in the *Survival Motor Neuron 1 (SMN1)* gene. SMN functions in the assembly of spliceosomal RNPs and is well conserved in many model systems including mouse, zebrafish, fruit fly, nematode, and fission yeast. Work in *Drosophila* has primarily focused on loss of SMN function during larval stages, primarily using null alleles or strong hypomorphs. A systematic analysis of SMA-related phenotypes in the context of moderate alleles that more closely mimic the genetics of SMA has not been performed in the fly, leading to debate over the validity and translational value of this model. We therefore examined fourteen *Drosophila* lines expressing SMA patient-derived missense mutations in *Smn*, with a focus on neuromuscular phenotypes in the adult stage. Animals were evaluated on the basis of organismal viability and longevity, locomotor function, neuromuscular junction structure, and muscle health. In all cases, we observed phenotypes similar to those of SMA patients, including progressive loss of adult motor function. The severity of these defects is variable, and forms a broad spectrum across the fourteen lines examined, recapitulating the full range of phenotypic severity observed in human SMA. This includes late-onset models of SMA, which have been difficult to produce in other model systems. The results provide direct evidence that SMA-related locomotor decline can be reproduced in the fly and support the use of patient-derived SMN missense mutations as a comprehensive system for modeling SMA.

## Introduction

Spinal Muscular Atrophy (SMA) is a neurodegenerative disease that primarily affects motor neurons in the anterior horn of the spinal cord and is a leading genetic cause of death among infants (Pearn, 1980). Symptoms involve muscle weakness that progressively worsens to the point of paralysis. The diaphragm becomes involved in later stages, leading to difficulty breathing and persistent respiratory infection that is a typical cause of death (Crawford, 2017). SMA has a broad range of severity; symptomatic onset can occur *in utero* in the most severe cases or in adulthood in the least severe. This spectrum has been subdivided into different “types” of SMA (Darras and Finkel, 2017; Talbot and Tizzano, 2017) based on age of onset: Type 0 (*in utero*), Type I (first 6 months), Type II (7 to 18 months), Type III (childhood onset after 18 months), and Type IV (adult onset). Although motor neurons are the most dramatically impacted cell-type in SMA, other tissues including the musculature, cardiovascular system, liver, pancreas, gastrointestinal tract, and immune system are also affected (Perez-Garcia *et al.*, 2017).

SMA is most commonly caused by reduced levels of the Survival Motor Neuron (SMN) protein which is encoded in humans by two genes, *SMN1* and *SMN2* (Lefebvre *et al.*, 1995). SMN protein is ubiquitously expressed and canonically functions in the assembly of spliceosomal snRNPs (Matera *et al.*, 2007; Matera and Wang, 2014; Gruss *et al.*, 2017). SMN is also reported to have functions related to RNA trafficking, translation, endocytosis, cytoskeletal maintenance, and cellular signaling (Raimer *et al.*, 2017; Singh *et al.*, 2017; Chaytow *et al.*, 2018; Price *et al.*, 2018). There is currently one FDA-approved SMA treatment: an antisense oligonucleotide (ASO) called nusinersen that increases SMN protein production from *SMN2* (Shorrock *et al.*, 2018). This therapeutic strategy prevents motor neuron dysfunction if treatment begins before or soon after symptomatic onset (Finkel *et al.*, 2017). In later-stage patients, ASO therapy halts degeneration but does not restore lost function (Mercuri *et al.*, 2018). Thus, these patients remain at high risk of complication and death despite receiving treatment.

The approved method of ASO delivery (intrathecal injection) treats only the central nervous system. While this is sufficient to prevent motor neuron dysfunction and early death, it appears likely that secondary symptoms will arise with age in peripheral systems of ASO treated patients (Hua *et al.*, 2015; Bowerman *et al.*, 2017). Therefore, there is an emerging need for SMA models that can be rapidly adapted to new research challenges. In this vein, *Drosophila melanogaster* is a highly useful model. In addition to the general advantages of the fly (low cost, rapid generation time, high quality genetic tools), this model system also has well characterized organ systems and cellular pathways that are highly relevant to the study of classical and emerging peripheral phenotypes in SMA.

*Drosophila* has a single *Smn* gene that is conserved, both in terms of protein sequence (Chan *et al.*, 2003) and molecular function (Rajendra *et al.*, 2007). To date, work in the fly has focused primarily on assessing the effects of strong *Smn* depletion on organismal development (Rajendra *et al.*, 2007; Shpargel *et al.*, 2009) or larval synapses and musculature (Chan *et al.*, 2003; Chang *et al.*, 2008), reviewed in (Grice *et al.*, 2013; Aquilina and Cauchi, 2018). Despite this body of work, the validity and translational value of the fly as a model for SMA continues to be called into question (Bowerman *et al.*, 2017; Iyer *et al.*, 2018). This appears to be due, at least in part, to the lack of a systematic and comprehensive analysis of SMA-related phenotypes at the organismal level. Here, we aim to fill this gap and more firmly establish the fly as a comprehensive system for the study of SMA by complementing the large body of existing work on the impact of *Smn* mutations on molecular, cellular, and neuromuscular phenotypes.

In addition, this work also fills a more general need in the field, namely, the modeling of intermediate forms of SMA. In many model systems, there are few effective models of SMA Types II and III (Burghes *et al.*, 2017; O’Hern *et al.*, 2017). In mouse, the many attempts to generate such models have almost invariably produced animals that are either severely affected, or completely unaffected in terms of neuromuscular phenotype (Le *et al.*, 2005; Osborne *et al.*, 2012). This is problematic not only for assessing intermediate forms of SMA, but also because nearly all severe SMA models exhibit developmental delays or arrest. This fact makes dissection of specific SMA-related phenotypes from general stress and death responses in an organism difficult and has complicated analysis of transcriptomic profiling of pre-mRNA splicing and neuromuscular development and function (Winkler *et al.*, 2005; McWhorter *et al.*, 2008; Hammond *et al.*, 2010; Garcia *et al.*, 2013; Garcia *et al.*, 2016).

Here, we present a set of fourteen SMA models that cover the full spectrum of SMA severity and, in many cases, circumvent problems of developmental delay. In these models, symptomatic onset occurs anywhere from early development to adulthood, suggesting that the platform effectively models the full spectrum of SMA Types. The larval stages appear particularly useful for examining pre- and early-onset SMA, as we observe reduced locomotor function in the absence of overt synaptic or muscular defects. Conversely, the adult stage is well suited for modeling onset and progression of SMA phenotypes, as we observed reduced lifespan and locomotor deficits at different times post-eclosion. Similar to human patients, loss of motor function in adult flies is progressive and displays early involvement of posterior limbs relative to anterior ones. These results provide evidence that this system is a valuable and effective tool for studying the full range of SMA pathogenesis.

## Materials and Methods

### Fly lines and husbandry

Patient mutation lines, *Smn^X7^*, *Smn^D^*, *Da-Gal4*, “C15” driver line: *elav(C155)-Gal4; Sca-Gal4; BG57-Gal4*. From Bloomington: *TRiP.JF02057* (Smn-RNAi #1, Stock #26288), *TRiP.HMC03832* (Smn-RNAi #2, Stock #55158).

To generate lines expressing *Smn* missense mutations, *Smn^X7^*/TM6B-GFP virgin females were crossed to *Smn^X7^*, *Smn^TG^*/TM6B-GFP males at 25°C. To reduce stress from overpopulation and/or competition from heterozygous siblings, crosses were performed on molasses plates with yeast paste and GFP negative, *Smn^X7^, Smn^TG^/Smn^X7^* larvae were sorted into vials containing molasses fly food during the second instar larval stage. Sorted larvae were raised at 25°C until the desired developmental stage was reached.

Experiments involving *UAS-Smn-RNAi* were carried out at 29°C to maximize expression from the Gal4/UAS system and, therefore, the degree of *Smn* knockdown. The one exception to this is the adult locomotion assay performed on Da-Gal4/Smn-RNAi #2. Raising these animals at 29°C dramatically reduces viability and is incompatible with survival to the adult stage. To circumvent this and produce viable adults we instead raised all animals for this experiment at 25°C. To maintain consistency across experiments, we also use molasses plates with yeast paste and subsequent sorting for all *Smn-RNAi* experiments.

### Viability Assays

To assess viability, we sorted 35-50 late second/early third instar larvae into vials containing standard molasses fly food and waited for them to complete development. After sufficient time had passed to allow for animals to complete development, we counted the total number of pupal cases in each vial and the number of empty pupal cases, which corresponds to the number of eclosed adults. We calculated % viability at both the pupal and adult stages by dividing these values by the number of initial larvae and then multiplying by 100 (pupal viability = (# total pupae/# initial larvae)*100, adult viability = (# empty pupal cases/# initial larvae)*100). Each vial is considered a biological replicate in respect to calculating averages and standard error. n-value represents the total number of larvae collected and assayed.

Determining the stage of pupal arrest involves an identical procedure to assessing pupal and adult viability with the exception of scoring the pupae. In this assay, we examined pupae under white light at 2X magnification and score them based on morphology as having arrested early in pupal development, in mid pupal development, late in pupal development, or as empty pupal cases (viable adults). These values were normalized to the total number of pupae.

### Locomotion Assays

Larval Locomotion: The locomotion assay used here was adapted from a previously published protocols (Brooks *et al.*, 2016). One to five larvae at either the early third or wandering third instar stage were placed onto the center of the locomotion stage (a large molasses plate) at room temperature. The stage was then placed in a recording chamber to control light and reflection and provide support for video recording. Once all larvae were actively crawling, movement was recorded as video for at least 1 minute and 10 seconds on an iPhone 6S at minimum zoom. Two video recordings were taken for each set of larvae. Locomotion videos were transferred to a PC and converted to raw video avi files in ffmpeg. Video length was trimmed to exactly 60 seconds by removing the first 2 seconds and final additional second to 1) create videos of standard duration and 2) eliminate from the analyzed frames small movements caused by starting and stopping the recording. Videos were opened and converted into binary images series in Fiji/ImageJ. The wrMTrck plugin for ImageJ (Husson *et al.*, 2013) was used to assess larvae size, larval body length, average speed of movement in pixels/second, and average speed normalized to larval size (body lengths/second).

Adult Locomotion: Adult flies of various ages were place in individual locomotion chambers consisting of 35 x 10 mm round tissue culture plates filled with 7.5 ml of clear agar to prevent movement in the Z-direction. Flies were given 5-6 hours to adjust to the new environment and then free moving walking behavior was recorded and analyzed in the same manner described for the larval locomotion assay.

### Immunostaining

Third instar larvae were dissected in HL3 saline with the following components (and concentrations): NaCl (70 mM), KCl (5 mM), MgCl2 (10 mM), NaHCO3 (10 mM), sucrose (115 mM = 3.9%), trehalose (4.2 mM = 0.16%), HEPES (5.0 mM = 0.12%), and CaCl_2_ (0.5 mM, unless otherwise indicated). Dissected samples were subsequently fixed with Bouin’s fixative (Ricca Chemical Company, Arlington, TX) for 3 minutes. Fixed samples were washed in 1X PBST using standard procedures, and incubated in primary antibodies at 4°C for overnight. This was followed by additional washes and a two-hour incubation in secondary antibody at room temperature. Staining was performed using the following primary antibodies: mouse anti-Synapsin (3C11) 1:1000 (concentrated antibody, Developmental Studies Hybridoma Bank, University of Iowa - DSHB); rabbit anti-Dlg 1:30,000 (Budnik *et al.*, 1996). The following fluorophore-conjugated antibodies were also used: Alexa-488 goat anti-rabbit 1:5000 and Alexa-546 donkey anti-mouse 1:5000 (both from ThermoFischer/Invitrogen Life Technologies, Eugene OR), and Alexa-647 goat anti-HRP (Jackson ImmunoResearch Laboratoreis, Inc., West Grove, PA). Larval preparations were mounted in Antifade media and imaged on a Leica TCS SP5 AOBS UV/spectral confocal laser-scanning system mounted on an inverted DM IRE2 microscope. Boutons were counted manually in Fiji/ImageJ in z-stacks were examined to insure that any boutons overlapping in the z-direction were accounted for. Muscle surface area was determined by outlining muscle 6 and 7 manually in Fiji/ImageJ. Muscle surface area is a rough measure, as rearing conditions and differences in muscle stretching can cause variability in the phenotype. Rearing conditions (both temperature and larval population within a vial) were carefully controlled and variability in stretching during dissection was minimized as much as possible.

### Longevity Assay

To assess longevity, newly eclosed adult flies were collected daily into vials containing molasses agar food. Animals were kept with no greater than 10 animals total to avoid stress from crowding and transferred to fresh vials every 2-3 days to avoid death due to non-optimal food conditions. The number of adults was recorded at the time of collection and once each week following collection until all animals had expired. Animals that escaped or were injured/killed during transfer were excluded from analysis.

### Collecting Partially Eclosed Adults

Animals were crossed and larvae collected as described above. As animals entered the pupal stage and began to either arrest and die or proceed through development. 2-3 days after pupal formation, viable pupae were transferred to an empty tissue culture dish for easy observation. Animals nearing the end of pupal development were observed for partial eclosion. Wild type flies eclose rapidly (within 1-10 seconds) under normal conditions. To be sure that Smn missense mutants were truly stuck in the partially eclosed state, we waited for 10 minutes following identification of potential partially-eclosed animals to see if they would complete eclosion. If they did not, we used fine forceps and microdissection scissors to remove partially eclosed animals from their pupal cases without damaging them. To control for the effects of possible stress during this assisted eclosion approach, we performed the same dissections/eclosion assist procedure on animals expressing wild type transgenic SMN that had completed pupal development but not yet eclosed.

### Scoring Melanotic Masses

Wandering third instar larvae were removed form vials, washed briefly in a room temperature water bath, dried, and placed on an agar plate under white light and 2X magnification. When melanotic masses were identified in a larvae, both the size of the largest mass (size score) and the total number of masses (mass score) were qualitatively determined. Size scoring used the following criteria: small masses range in size from barely visible specks to smooth round dots with a diameter no more than 1/10^th^ the width of the larva; medium masses range from anything larger than a small mass to those with a diameter up to 1/3 the larval width; large masses are any with a diameter greater than or equal to 1/3 the larval width. Larvae were manipulated to allow for observation of all sides/regions; observation was performed for at least 20 seconds in all cases.

### Statistical analysis

All statistical analyses were performed using GraphPad Prism 7. For all data except longevity curves, p-values are from one-way ANOVA with a Dunnet correction for multiple comparisons. Statistical significance for longevity curves was determined by the Logrank/Mantel-Cox test. Details for the statistical analysis used in each figure panel are described in figure legends and/or shown in Supplemental Table 1.

## Results

To maximize the range of phenotypic severity examined in this study, *Drosophila* lines carrying transgenic insertions of fourteen different *Smn* alleles were examined, in addition to the Oregon R (OR) wild type lab strain. The inserted transgenes encode a wild type *Smn* allele (WT) along with thirteen alleles for non-synonymous *Smn* point mutations. Each mutation in this allelic series produces SMN protein with a single residue change homologous to those of human SMA patients bearing small *SMN1* mutations (Supp Table 2). In this work, we will refer to these transgenic lines by the amino acid substitution produced in the fly SMN protein (D20V, for example). A single copy of each missense mutation allele is expressed in an *Smn* null background (see Materials and Methods for full genotype). Thus the transgenic product is the sole form of SMN present in the animals after the maternal SMN contribution is fully depleted at the end of the second instar larval stage. For some Smn mutations, we have also generated stable, self-replicating lines with animals carrying the Smn transgene and two transheterozygous null mutations of the endogenous *Smn* gene. These flies lack maternally contributed wild type Smn at all stages of development.

It should be noted that expression of each of these transgenic *Smn* alleles is driven by the native 5’ and 3’ *Smn* flanking regions and inserted at an identical attP site on the third chromosome to standardize potential position effects. This genomic arrangement is reported to drive transgenic *Smn* expression at lower levels than the endogenous *Smn* gene (Praveen *et al.*, 2012). The *Smn*^WT^ transgenic line serves both as a control for this lower level of SMN expression and, in some assays described here, displays mild phenotypic changes relative to the wild type lab strain Oregon R (OR). All but one of these *Smn* missense lines has been previously reported (Praveen *et al.*, 2014). This work reports that these *Smn* alleles causes variable degrees of developmental arrest and reduced viability, suggesting that they could be useful in recapitulating the spectrum of SMA severity seen in human patients. Beyond this observation, the authors take a molecular approach and focus their analysis on the impact of these *Smn* point mutations on protein interactions within the SMN complex. Only the most severe alleles in the series (G206S, Y203C, and M194R) were assayed for reduced locomotor behavior as early third instar larvae. Beyond this, no phenotypic analysis of SMA-related phenotypes has previously been assessed in this allelic series.

Here, we fully characterize these *Smn* alleles in the context of organismal SMA-related phenotypes. We also report on an additional *Smn* allele based on a recently reported SMA mutation that produces a tyrosine to cysteine change at Y277 in the human (Y208 in *Drosophila)* SMN protein (Supplemental Table 2).

### *Smn* missense mutations reduce viability across development

We first sought to replicate the previously reported viability defects for each of these 14 lines at the pupal and adult stages (Fig. 1A). Consistent with previously published observations, our experiments produced a broad range of viability phenotypes. The most severe phenotype is full developmental arrest at the 2^nd^ to 3^rd^ instar transition, which was observed for the G206S, M194R, and Y203C alleles. Animals carrying any of the other eleven *Smn* alleles reach pupation at moderate to normal levels, and all but one of them (V72G) produce viable adults to some degree (Fig 1A). The adult viability defects form a continuum that encompasses phenotypes from complete lethality (V72G) to nearly normal levels of viability (D20V and WT). This continuum is reminiscent of the spectrum of disease severity seen in SMA patients, indicating that this set of missense mutations has the potential to model a broad range of SMA Types.

**Figure 1.**
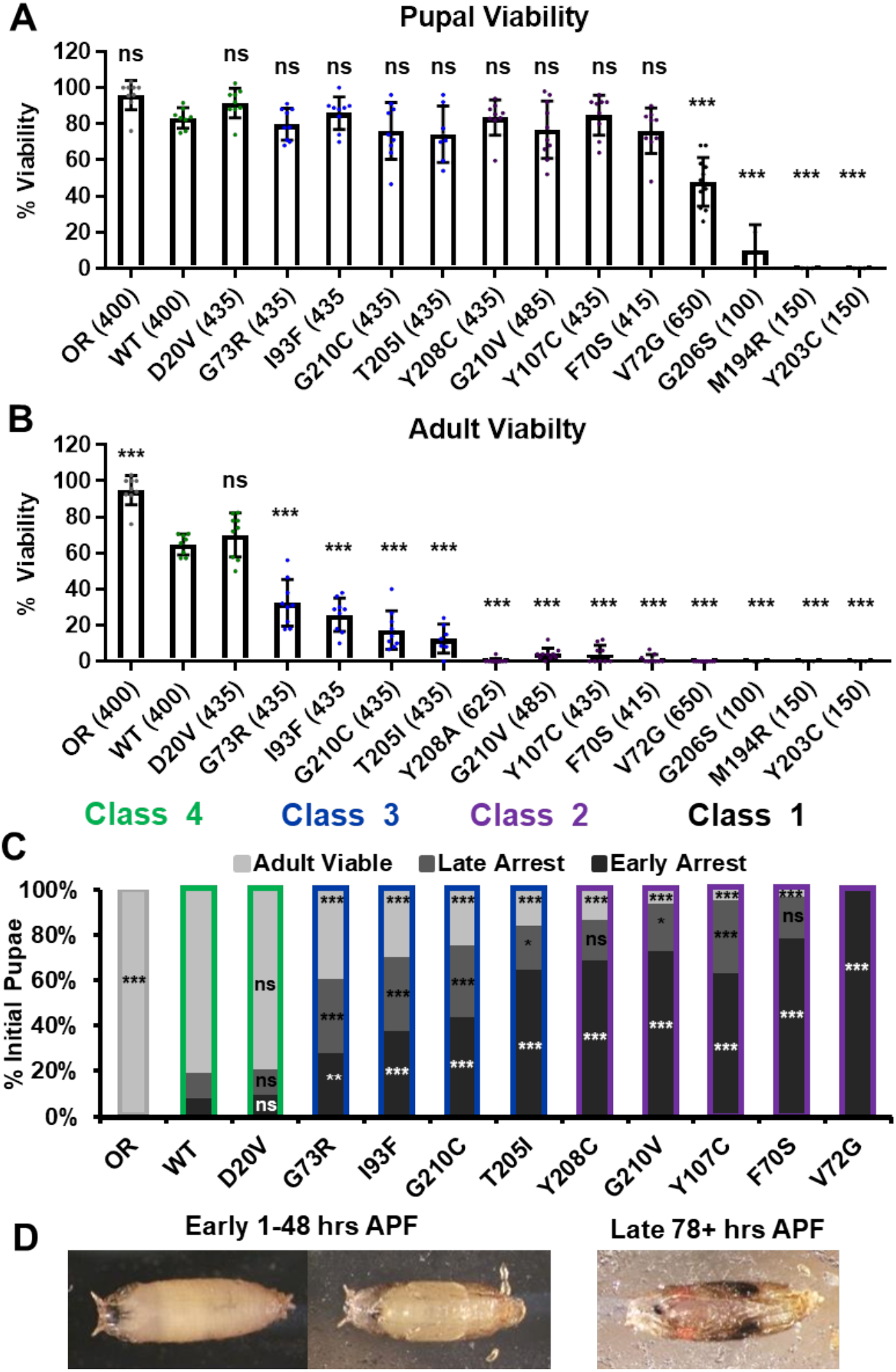
*Smn* missense mutations produce a spectrum of viability defects. **A and B)** Developmental viability of animals expressing *Smn* missense mutations at the pupal stage (A) and the adult stage (B) *%* Viability is the proportion of animals that survive to the pupal/adult stage relative to the number of larvae initially collected. **C)** Viability of animals carrying *Smn* missense mutations through pupal development. In the case, ns indicates a p-value of 0.99 and *** indicates a p-value of exactly 0.0001 **D)** Representative images of the pupal stages assessed in C. **Data:** Bars show average. Error bars show standard error. Data points represent biological replicates of 35-50 animals each, n-values (shown as numbers in parentheses next to genotypes) reflect the number of individual animals counted. n-values are the same for panels A-C. **Statistical analysis:** Values above the data indicate significance vs WT from one-way ANOVA using the Dunnet correction for multiple comparisons. ns: not significant (p>0.05), * p<0.05 **p<0.01, ***p<0.001

A majority of these *Smn* alleles arrest during the pupal stage of development. To further explore this phenotype, we examined the timing of death during pupariation for each allele (Fig 1B,C). The pupal cases of developing *Drosophila* are transparent, so approximate staging can be performed visually using intact animals (Fig 1C). We observed arrest and death both early and late during pupal development for animals carrying all alleles except V72G, which causes arrest very shortly after pupariation (Fig 1B,C). Very few animals of any line arrested during the middle stages of pupation. Higher frequencies of arrest early in pupal development were observed for alleles that produce fewer viable adults.

Given the large number of individual genotypes described here, we organized these *Smn* alleles into four phenotypic classes based on gross viability phenotypes. Class I is the most severe, causing larval-stage arrest, and is comprised of G206S, M194R, and Y203C. Class II is comprised of the five most severe alleles that reach pupation: V72G, F70S, Y107C, G210V, and Y208C. A small fraction (2-11%) of animals carrying Class II alleles eclose as adults. Class III alleles are more moderately affected, with ecolsion frequencies ranging from 20-45%. This class includes the T205I, G210C, I93F, and G73R alleles. Class IV includes the WT and D20V alleles, which are the least affected, displaying a moderate 20-30% decrease in adult viability relative to the Oregon R strain. Throughout this manuscript, data for the *Smn* alleles are arranged in order of severity, and the color scheme for each Class is maintained for ease of reference.

### *Smn* missense mutations support viability when wild type SMN protein is completely absent

In combination with previous observations, the range of severities observed in our viability assays suggests that most *Smn* missense mutations retain at least partial function. This conclusion is complicated by the presence of maternally contributed SMN that is present during embryonic and early larval development. We therefore attempted to generate stable fly lines expressing only *Smn* missense mutations. To do this, we sought to generate flies carrying two different *Smn* null alleles (*Smn^X7^* and *Smn^D^*) as well as one copy of each transgenic *Smn* allele (*Smn^TG^*) for Class III and Class IV alleles. In all cases, adults of the desired genotype (*Smn^TG^, Smn^X7^/Smn^D^*) were viable and sufficiently fertile to produce self-sustaining stocks. In later generations, animals in these stocks animals can carry either one or two copies of the Smn transgene and completely lack wild type SMN in all stages of life and development.

Having created these stocks, we assessed the functionality of *Smn* missense mutations by measuring developmental viability (Supp Fig 1). We found that, in all cases, expression of Class III and Class IV mutations is sufficient for robust pupal viability (Supp Fig 1A) and adult viability ranging from the moderate to wild type levels (Supp Fig 1B). This result definitively demonstrates that *Smn* missense mutations are functional in the absence of any wild type SMN protein.

### *Smn* missense mutations cause defects in larval locomotion but not NMJ structure

Manipulations that strongly reduce SMN levels are known to cause larval locomotion defects. It is not clear, however, if the moderate reduction of SMN function modeled by *Smn* missense alleles has the same functional impact as strong reduction of wild type SMN. To address this, we performed larval locomotion assays on all lines at the early third instar stage (72 hours post-egg lay) and in wandering third instar larvae for the eleven lines that reach this stage (Fig 2).

**Figure 2.**
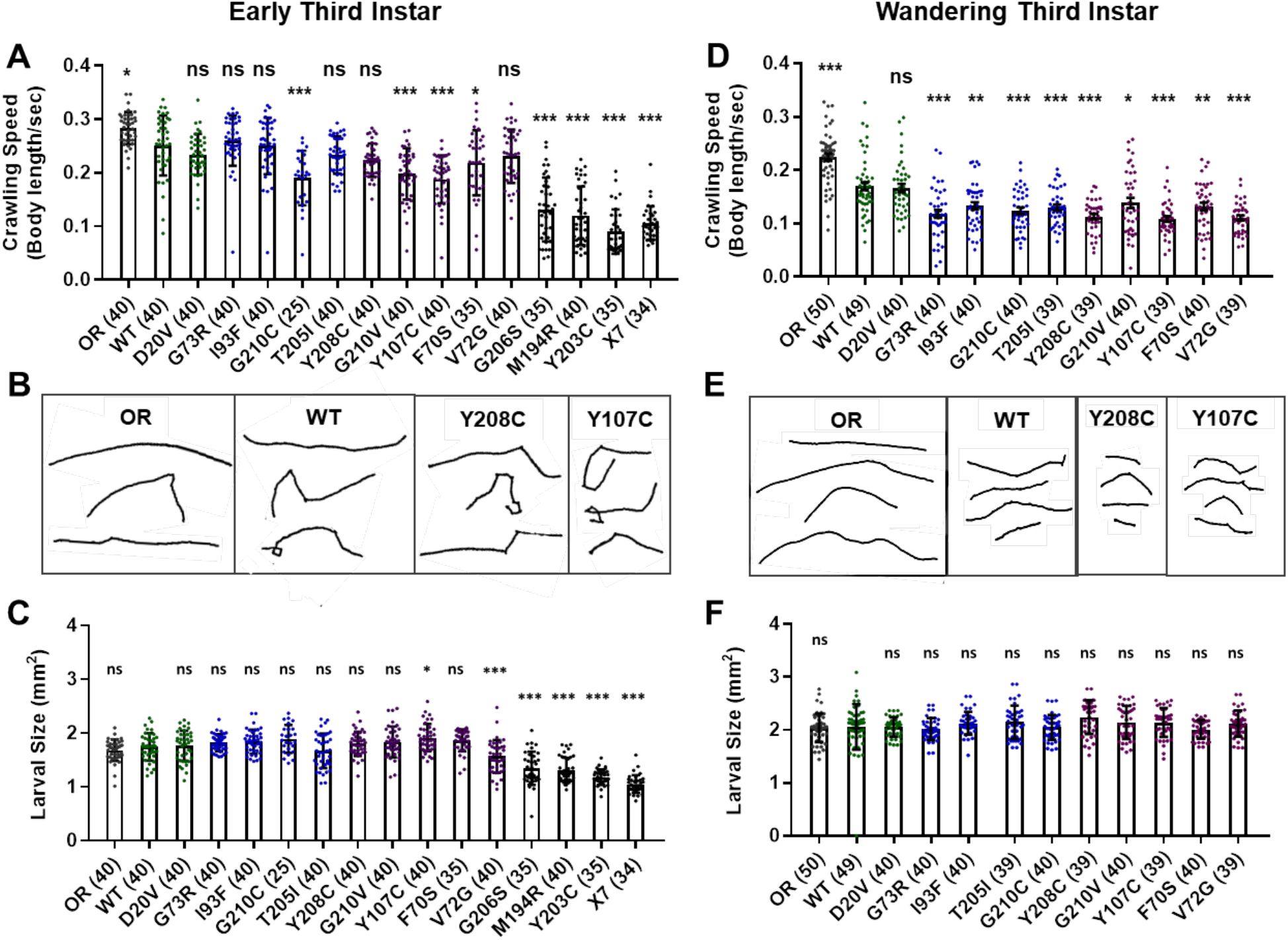
*Smn* missense mutations or knockdown reduces larval locomotion. **A)** Larval crawling speed, measured in body lengths/second, in early third instar larvae. **B)** Representative traces for the data shown in A. **C)** Larval size for the same animals measured in the locomotion assay in A. **D)** Larval crawling speed, measured in body lengths per second, in wandering third instar larvae. **E)** Representative traces for the data shown in D. **F)** Larval size for the same animals measured in the locomotion assay in D. Data Bars show averages, points represent individual larvae, error bars represent standard error, n-values (# individual larvae) are show in parentheses adjacent to genotype on x-axis. Statistical significance was determined by ANOVA (ns = not significant, *p<0.05, ** p<0.01, ***p<0.001)

Larvae from all lines display reduced locomotion relative to Oregon R at the early third instar stage with the exception of G73R (Fig 2A,B). All three Class I alleles show a further decrease in crawling speed relative to the wild type transgenic line, as do the G210C, G210V, F70S, and Y107C lines. The rest of the mutant lines display no significant difference from the wild type transgenic control, suggesting that the mildly hypomorphic nature of these alleles is driving their locomotor defects (Fig 2A,B).

By the wandering third instar stage, all surviving lines exhibit locomotion defects. The Class IV alleles show a mild but statistically significant reduction in crawling speed relative to the Oregon R wild type train (Fig 2D). These two Class IV *Smn* alleles (WT and D20V) are not significantly different from one another, whereas all Class II and three alleles show a significant reduction in locomotion relative to both OR and the WT transgenic lines (Fig 2D,E). These differences are not due to overall changes in larval size, as this measure is unchanged in the wandering third instar larvae carrying transgenic *Smn* alleles (Fig 2F) and is controlled for within the analysis, by measuring locomotion in terms of body lengths per second (BLPS).

Given these locomotor deficits and previous reports of reduced NMJ size in the case of strong loss of SMN, we next immunostained and counted boutons at the muscle 6/7 NMJ in abdominal segment A2 for all lines that reach the wandering third instar stage. Surprisingly, we did not observe a dramatic change in bouton number for any of these alleles. All lines showed a slight but non-significant decrease in bouton number relative to Oregon R, with the exception of the I93F allele, which does reach statistical significance (Supp Fig 2 A,B). These findings suggest that the locomotion defects observed in these larvae are likely due to functional changes in motor neurons or in other upstream central neurons such as interneurons. We also assessed the combined surface area of muscle 6/7 in segment A2 to look for signs of muscle degeneration. At the wandering third instar, we observed no changes in muscle area for any *Smn* allele (Supp Fig 2C). This indicates that the body wall muscles are not actively experiencing atrophy and that these larvae are in the early stages of symptomatic onset.

### RNAi-mediated *Smn* knockdown phenocopies *Smn* missense alleles

We next assessed potential tissue-specific roles of SMN in the observed phenotypes. To address this question, we used the *Gal4/UAS* system to drive two different constructs that produce shRNAs targeting RNAi-based knockdown of *Smn* under control of a *UAS* enhancer element (Perkins *et al.*, 2015). These lines are *Smn-RNAi^JF02057^* and *Smn-RNAi^HMC03832^* and are referred to here as Smn-RNAi #1 and Smn-RNAi #2, respectively. Expression of these *UAS-RNAi* constructs is driven either ubiquitously with *Da-Gal4 (Daughterless)* or concurrently in both neurons and muscles using the ‘C15’ driver line (Brusich *et al.*, 2015), which contains three Gal4 constructs: two pan-neuronal drivers, *elav(C155)-Gal4* and *Scaberous-Gal4*, and one muscle specific driver, *BG57-Gal4*. Crossing the C15 driver with the Smn-RNAi responder lines allows us to simultaneously deplete SMN in both muscles and neurons, the two cell types most closely linked to SMA pathology.

We first used these RNAi lines to asses pupal and adult viability (Fig 3A,B). Similar to the findings with the *Smn* allelic series, both ubiquitous and neuromuscular *Smn* knockdown caused a significant reduction in viability (Fig 3A). Ubiquitous knockdown causes an effect comparable to that of the V72G lines, strongly reducing pupal lethality and displaying complete lethality at the adult stage. In contrast, neuromuscular knockdown of *Smn* phenocopies the moderate Class III alleles, displaying fairly normal pupal viability and a moderate decline in adult viability (Fig 3A). These data suggest that neuronal and muscle tissues contribute to the viability defects observed following ubiquitous *Smn* knockdown and in animals ubiquitously expressing mutant *Smn* alleles.

**Figure 3.**
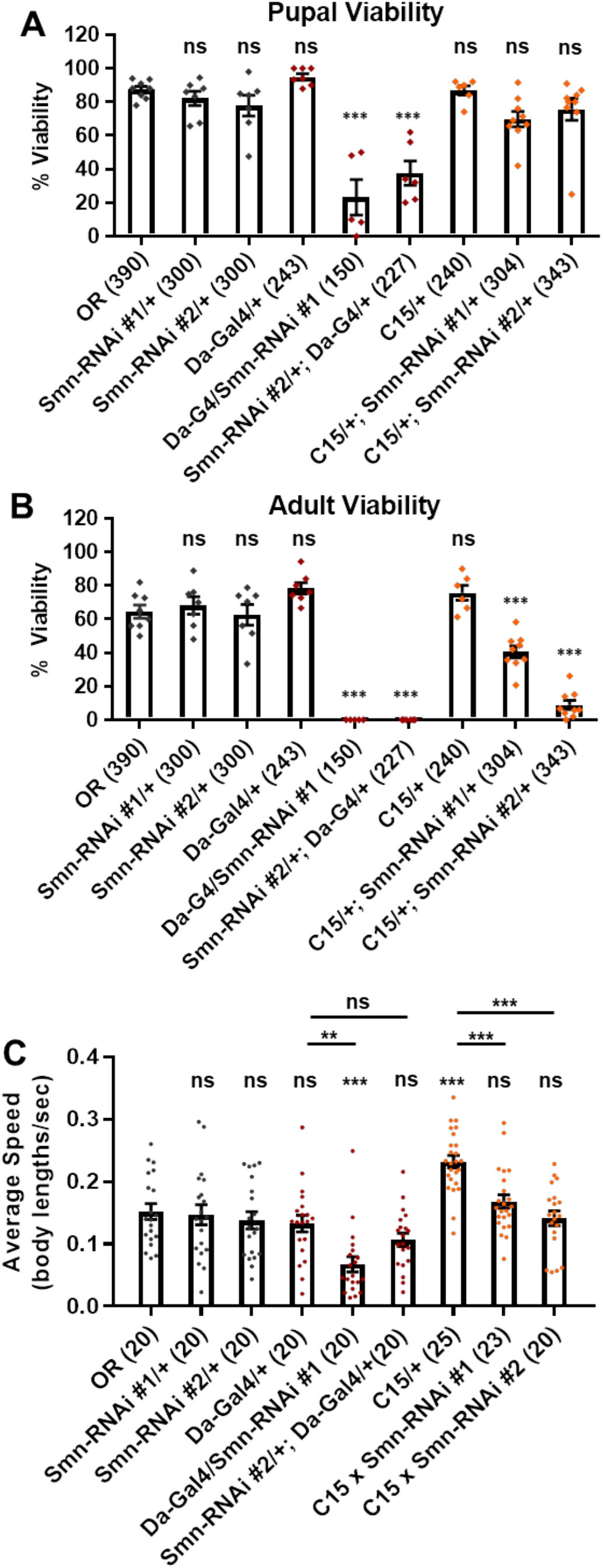
Neuromuscular *Smn* knockdown reduces viability and larval locomotion. **A and B)** Developmental viability at the pupal stage (A) and adult stage (B) for animals expressing *Smn* missense mutations. % Viability is the proportion of animals the survive to the pupal/adult stage relative to the number of larvae initially collected. **C)** Larval crawling speed, measured in body lengths/second, in early third instar larvae. **Data:** Bars show average. Error bars show standard error. For A and B, data points represent biological replicates of 30-50 animals each, n-values (shown in parentheses next to genotypes) reflect the number of individual animals counted. For C, data points and n-values (shown in parentheses next to genotypes) reflect the number of individual animals assayed. **Statistical analysis:** Values above the data indicate significance vs WT from one-way ANOVA using the Dunnet correction for multiple comparisons. ns: not significant (p>0.05), * p<0.05 **p<0.01 ***p<0.001

We also examined locomotion in the context of *Smn* knockdown. Both ubiquitous and neuromuscular knockdown of *Smn* negatively impacted larval locomotion (Fig 3C). In the context of ubiquitous knockdown, the presence of the *Gal4* element alone did not impact the baseline locomotor phenotype. In contrast, *Gal4* expression in neuromuscular tissues from the C15 line caused a significant increase in crawling speed relative to the OR control (Fig 3C). Relative to this increase, expressing Smn-RNAi using the C15 driver line caused a significant decrease in locomotor speed (Fig 3C). A similar decrease in larval velocity was also observed in the context of ubiquitous *Smn* knockdown. Specifically, ubiquitous knockdown with Smn-RNA #1 causes a significant reduction in crawling speed (Fig 3C). Unlike the lines expressing *Smn* missense alleles, ubiquitous knockdown also caused a significant reduction in the size of the muscle 6/7 neuromuscular junction relative to the Oregon R wild type strain (Supp Fig 2A).

Overall, examination of SMA-related phenotypes in larvae reveals a spectrum of severities in the context of developmental viability and reveals a moderate reduction in locomotor behavior. However, we did not observe changes to larval NMJ structure or muscle size (Supp Fig 2). This is likely due to the relatively mild to intermediate nature of the *Smn* mutations examined here, the effects of maternally contributed SMN, and the short duration of the larval stages. In the context of our SMA models, there is likely not enough time for degenerative phenotypes to manifest in the larval stages. Therefore, to examine potential progressive degenerative phenotypes, we turned to the adult stages.

### *Smn* missense mutations and neuromuscular knockdown reduce adult longevity

Ten lines expressing transgenic *Smn* alleles produce viable adults to varying degrees (Fig 1A). We first used these adults to assess longevity, with the expectation that *Smn* mutations would shorten lifespan. Animals of the control Oregon R strain lived as long as 12 weeks, with 90% of individuals dying by 11 weeks of age (Fig 4A-D, gray line). The line expressing transgenic *Smn* (WT) is moderately hypomorphic and shows a significant decrease in longevity as compared to OR, with increased death occurring by week 4 in the case of WT (Fig 4A-C).

**Figure 4.**
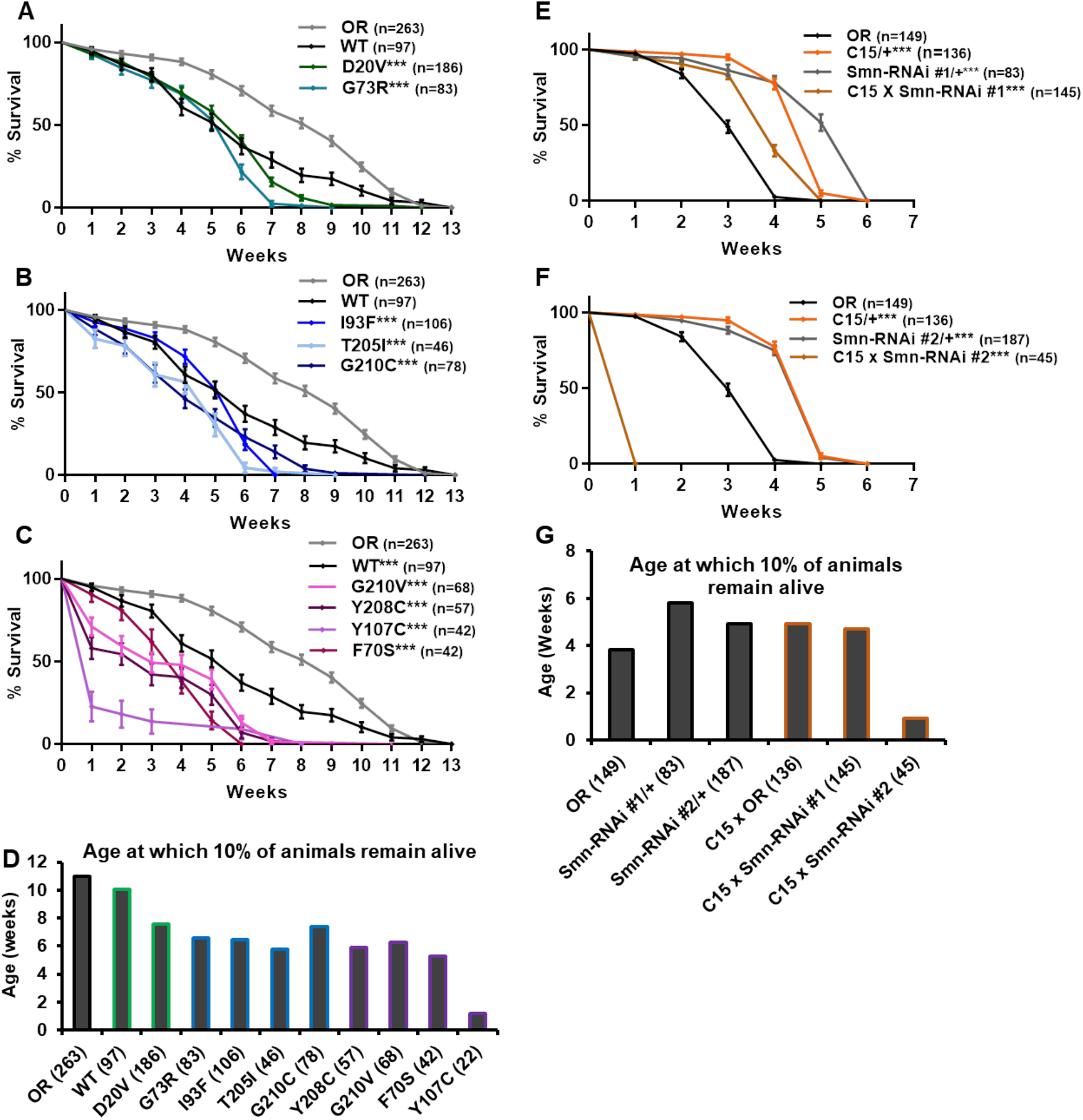
*Smn* missense mutations and *Smn* knockdown reduce lifespan. **A-C)** Survival curves for adult flies expressing *Smn* missense mutations. Data is split into three graphs for visual clarity. Colors correspond to phenotypic Class. **D)** Lifespan measured as the age at which 10% of animals remain alive for each missense mutation. **E and F)** Survival curves for adult flies expressing Smn-RNAi in all neurons and muscles. **G)** Lifespan measured as the age at which 10% of animals remain alive for each RNAi condition and control. Shown next to genotypes, n-value (# of individual flies), *** indicates p<0.0001 by Chi square analysis using the logrank rank/Mantel-Cox test.

All other lines expressing missense *Smn* alleles show a further, statistically significant reduction in longevity relative to WT (Fig 4A-D). The Class IV mutation, D20V, begins to display impaired longevity relative to WT beginning at week 7 and 90% of animals carrying this mutation are deceased by 7.6 weeks of age (Fig 4A, D). The decrease in longevity is more pronounced for the Class III mutations (G73R (Fig 4A), I93F, T205I, and G210C (Fig 4B), which begin to deviate from WT between 3-6 weeks of age (Fig 4B,C) and reach 90% death by 6-7 weeks (Fig 4D). The Class II mutations (G210V, Y208C, Y107C, and F70S) are the most severely impacted, with significant death occurring as early as the first week of life (Fig 4C,D).

We next sought to assess the possibility that the reduced longevity observed for missense *Smn* lines is related to neuromuscular health. To do so, we again turned to RNAi-mediated *Smn* knockdown in the neuromusculature. The C15 Gal4 driver line was used to express each of the two *UAS-Smn-RNAi* constructs in adult flies. To maximize shRNA expression, flies were raised at 29°C which increases the efficiency of the yeast-derived Gal4 protein as a transcriptional activator. Under these conditions, the OR strain displays reduced longevity/lifespan relative to other controls such as the Gal4 alone (C15/+) or the Smn-RNAi constructs alone (Smn-RNAi/+) (Fig 4E-G). Neuromuscular expression of Smn-RNAi #1 moderately affected longevity relative to the C15/+ and Smn-RNAi #1/+ controls, although they outlived the OR controls (Fig 4E,G). The effect with Smn-RNAi #2 was much stronger, with all animals expiring within the first week of life (Fig 4F,G). These findings suggest that the longevity defects observed in the case of missense *Smn* alleles is due, at least in part, to loss of *SMN* function in neurons and muscles.

### *Smn* missense alleles and neuromuscular knockdown reduce adult locomotion

To more directly assess neuromuscular function in adult flies we performed an adult locomotion assay on free moving animals in a circular chamber with restricted vertical space to prevent flight. Flies were collected and assayed within 24-36 hours after eclosing (Fig 5) and after aging for 5 and 6 weeks in certain cases (Fig 6). At one day of age, both Class IV alleles (WT and D20V) and two of the Class III alleles (G73R and I93F) showed robust locomotor behavior. These animals display walking speeds that are slightly faster than that of the Oregon R wild type strain and are not significantly different from one another (Fig 5A, B). The remaining Class III alleles (T205I and G210C) showed a moderate but significant decrease relative to the WT transgenic line. The Class II alleles showed robust and significant impairment in their locomotor behavior relative to both WT and Oregon R (Fig 5A,B).

**Figure 5.**
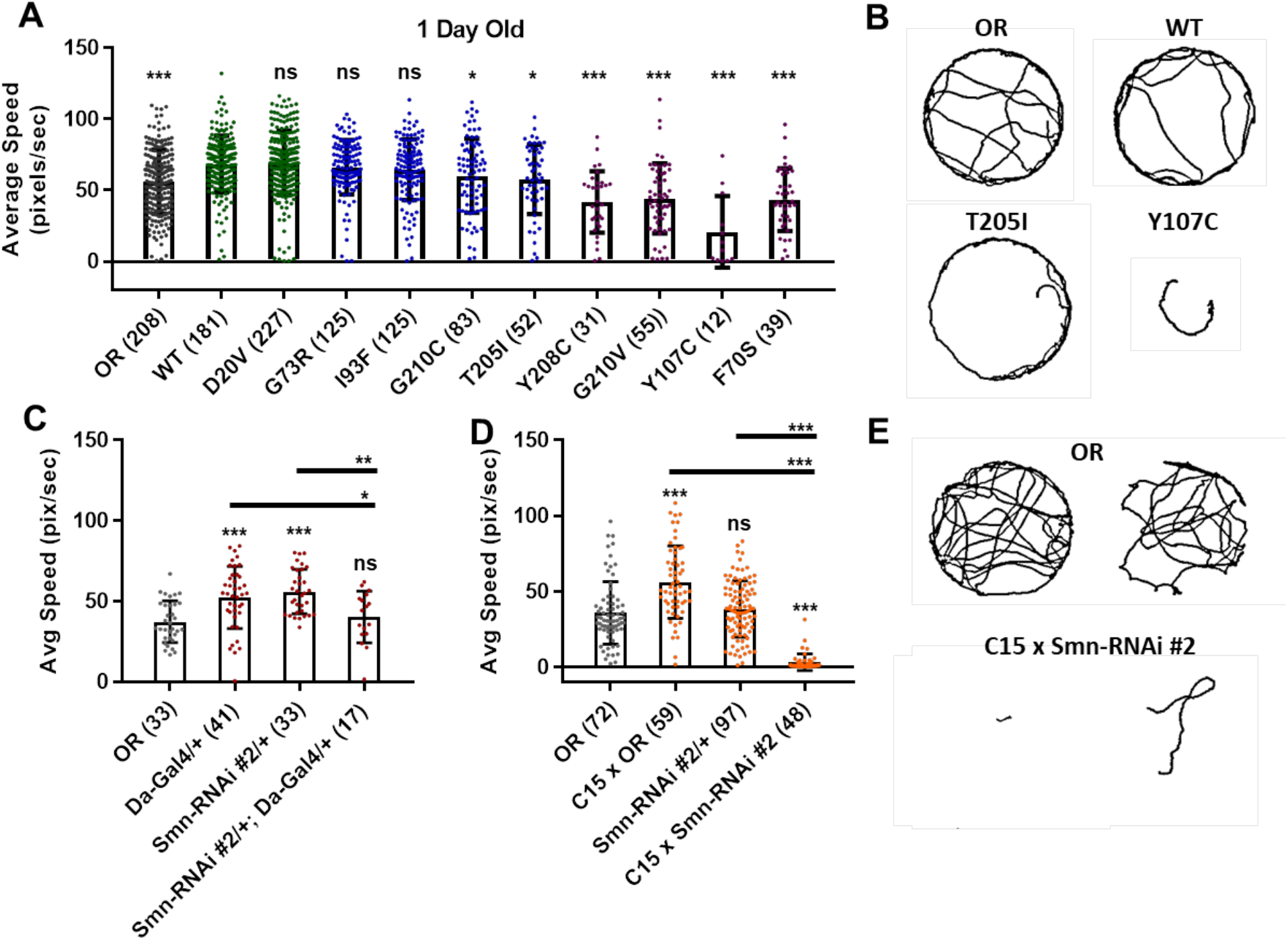
*Smn* missense mutations or knockdown reduces free moving adult locomotion. **A)** Adult walking speed, measured in pixels/second, in adults one day after eclosion for animals expressing *Smn* missense mutations. **B)** Representative traces for the data shown in A. **C and D)** Adult waking speed, measured in pixels per second, in adults one day after eclosion for animals expressing *Smn-RNAi* either ubiquitously with the Da-Gal4 driver (C) or in both neurons and muscle using the C15 driver line (D). **E)** Representative traces for the data shown in D. **Data:** Bars show average. Error bars show standard error. Data points and n-values (shown in parentheses next to genotypes) reflect the number of individual animals assayed. **Statistical analysis:** Values above the data indicate significance vs WT from one-way ANOVA using the Dunnet correction for multiple comparisons. ns: not significant (p>0.05), * p<0.05 **p<0.01 ***p<0.001

Similar effects were observed for both ubiquitous and neuromuscular knockdown of *Smn* (Fig 5C-E). As was the case for larval locomotion, Gal4 expression alone leads to increased adult walking speeds. Relative to Gal4 controls, ubiquitous *Smn* knockdown using the Smn-RNAi #2 line produces a moderate but significant reduction in adult walking speed (Fig 5C). Neuromuscular *Smn* knockdown with Smn-RNAi #2 also reduces walking speed relative to both the C15 Gal4 line control and the Oregon R wild type strain. In this case the reduction was dramatic, as many animals remained essentially stationary during the assay (Fig 5D,E). This finding suggests that the locomotor defects observed in animals expressing mutant *Smn* alleles are due to dysfunction in neuronal and/or muscular tissues. Collectively, these data suggest that the adult stage can be used to model SMA-related phenotypes in the fly.

### Class IV and Class III Smn missense alleles are models for late-onset SMA

Three missense *Smn* alleles (D20V, G73R, and I93F) displayed robust adult locomotor behavior at one day of age (Fig 5A,B). In contrast, these lines also show reduced longevity as adults (Fig 4), mildly reduced viability (Fig 1), and impaired locomotion as larvae (Fig 2). These results indicated that these mutations have a relatively mild impact on *SMN* function and led us to hypothesize that these lines might be useful for modeling late-onset SMA. To test this, we aged flies carrying each of these three mutations as well as animals expressing the WT *Smn* transgene and assayed adult walking behavior weekly (Fig 6A). By 5 weeks of age, flies expressing the D20V, G73R, or I93F mutant *Smn* all showed a significant reduction in walking speed relative to the animals expressing the WT *Smn* transgene (Fig 6A,B,D). This decline persisted into the 6^th^ week of life (Fig 6A,C,D), but could not be effectively assessed beyond that point due to the limited number of animals that survive into the 7^th^ week of life. Interestingly, this decline in locomotor function occurs 1-2 weeks prior to the time at which most of these flies began to die in the longevity assay (Fig 4), consistent with a causative link between the locomotor deficit and death. Based on this late-onset locomotor dysfunction, we conclude that these lines represent the first models of late-onset SMA that involve onset of dramatic impairment of locomotor function at an advanced age.

**Figure 6.**
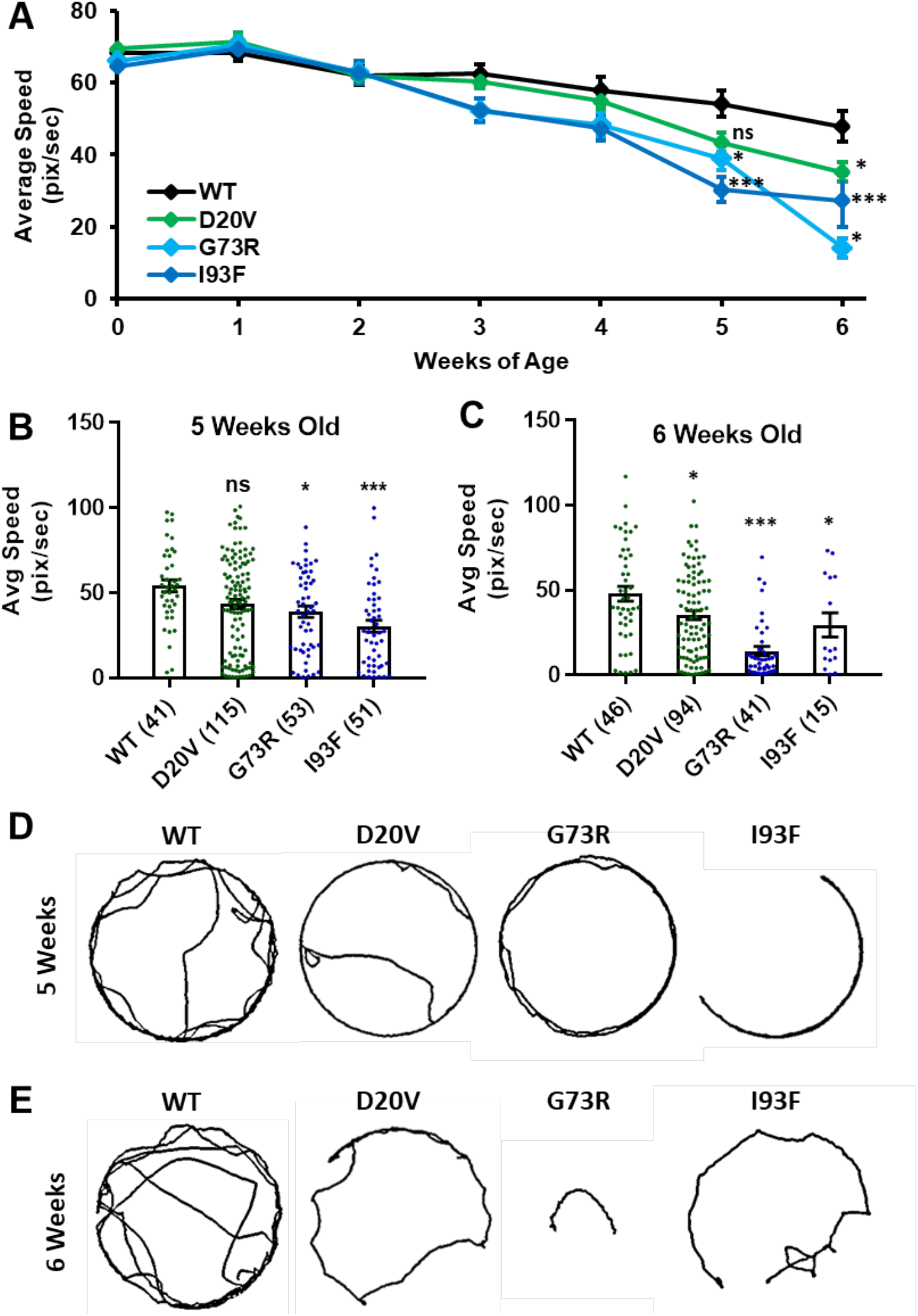
Animals expressing mild *Smn* missense mutations develop late-onset locomotor defects as adults. **A)** Adult walking speed, measured in pixels/second, in adults ranging from one day of age (0 weeks) through 6 weeks of age for animals expressing the WT transgene or the D20v, G73R, I93F *Smn* missense mutations. **B and C)** Adult walking speed (pixels/second) for the same genotypes shown in A at either 5 weeks or age (B) or 6 weeks of age (C). **D)** Representative traces for the data shown in B **E)** Representative traces for the data shown in C. **Data:** In A, points show averages in B and C, bars show average. Error bars show standard error in all cases. Data points and n-values (shown in parentheses next to genotypes) reflect the number of individual animals assayed. **Statistical analysis:** Values above the data indicate significance vs WT from one-way ANOVA using the Dunnet correction for multiple comparisons. ns: not significant (p>0.05), * p<0.05 **p<0.01 ***p<0.001

### Smn mutant alleles and knockdown cause progressive loss of motor function

We analyzed the more severe Class II lines for progressive loss of motor function. We noticed that certain animals failed to fully eclose from their pupal cases and ultimately died (Supp Fig 3A,B). This phenotype was most prevalent for the intermediate Class III mutations, and was less frequently observed for the more severe Class II mutations. This inverse relationship between allele severity and incidence of partial eclosion is likely due to the fact that most animals carrying Class II mutations arrest prior to reaching these later stages of development (Fig 1A,B).

We hypothesized that partial eclosion occurs in part due to severe muscle weakness in a subset of animals expressing missense *Smn*. To test this, we selected two lines, Y208C and G210V, which are both phenotypically severe in other assays yet produce enough late-stage animals to make analysis feasible. We monitored pupae for these lines on an hourly basis to identify animals that were unable to escape their pupal cases. Once identified, forceps and microdissection scissors were used to help these animals complete eclosion (“assisted eclosion”). We first put these animals through the adult locomotion assay and found that they had very low walking speeds and in many cases did not move at all (Supp Fig 3C). Additionally, the lifespan of these animals was dramatically reduced relative to both wild type and normally eclosing adults of the same genotype (Supp Fig 3D, Fig 4C,D). All animals examined died within 6 days of assisted eclosion (Supp Fig 3D). To ensure that our assisted eclosion techniques were not harming the animals or impairing their locomotor function or survival, we performed the same procedure on *Smn^WT^* (WT) pharate adults. Analysis of lifespan and locomotor function of these animals was included for control purposes. As expected, the WT animals were unaffected by the manipulation (Supp Fig 3C,D).

Similar phenotypes were also observed for animals expressing neuromuscular knockdown of *Smn* with Smn-RNAi #2. These animals are able to eclose without assistance but display severe adult locomotor dysfunction in the adult walking assay one day after eclosion (Fig 5F,G). Neuromuscular *Smn* knockdown also leads to dramatically reduced lifespan almost identical to that seen for the partially eclosed animals examined, with all animals dying within 6 days of life (Supp Fig 3A).

Although walking speed was near zero for all of the mutant lines discussed above, qualitative observation suggested that these animals were not fully paralyzed and retained some degree of motor control over their legs at the time of assisted eclosion. To determine if motor control deteriorated over time in these animals, we turned to qualitative observation and scoring of locomotor function in a manner similar to clinical analysis of human patients. For 10-30 individual animals per genotype, we monitored leg function under the stereoscope for 1-2 minutes and assigned a leg score for each of the three leg pairs. Scoring was performed on a scale of 0 to 10, with complete paralysis scored as 0 and normal function scored as 10. Scoring was performed every 4-8 hours for the first 12 hours of life, followed by observation every 24 hours until death occurred. Using this method, we determined that partially eclosed flies expressing G210V and Y208C mutant *Smn* experience progressive loss of leg function over time (Fig 7A-C). This is the first report of progressive loss of motor function in any fly model of SMA.

**Figure 7.**
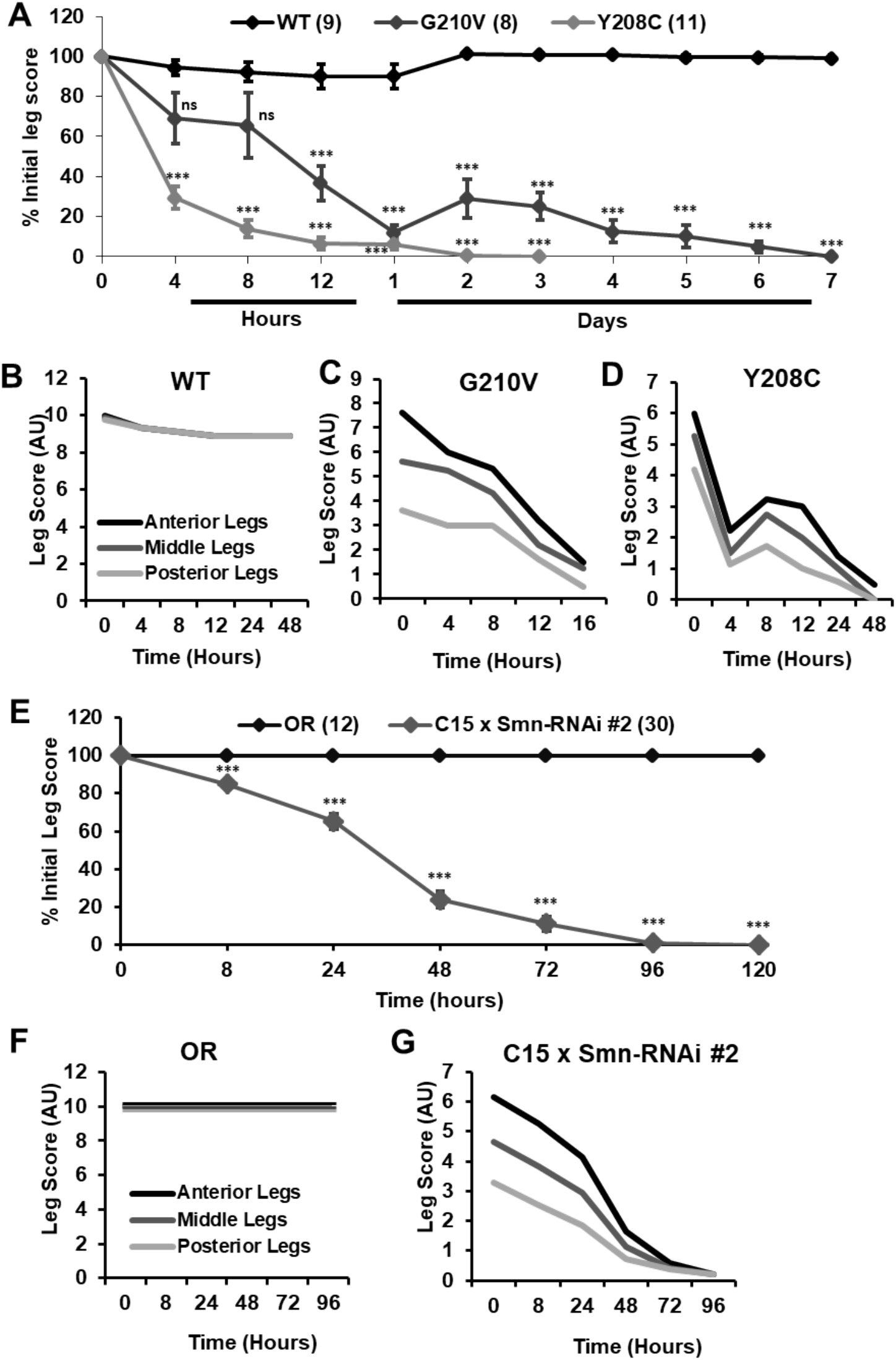
Progressive loss of motor function is observed in the legs of partially eclosed *Smn* missense mutants and in animals expressing neuromuscular Smn knockdown. **A** Qualitative leg function scores (average or all three leg pairs) over time for partially eclosed animals expressing the wild type *Smn* transgene (n=11) or the Y208C (n=10) or G210V (n=7) *Smn* missense mutations. **B-D)** Qualitative leg function scores for each leg pair over time for same animals as in A. **E)** Qualitative leg function scores (average or all three leg pairs) over time for animals expressing neuromuscular *Smn* knockdown. **F and G)** Qualitative leg function scores for each leg pair over time for same animals as in E.

Interestingly, individual leg pairs were differentially affected at the time of assisted eclosion. The most anterior pair initially presented with mild to moderate impairment, whereas the most posterior pair was more severely impaired. The function of all three leg pairs decreases over time, and the defects in the anterior and posterior limbs become more similar over time (Fig 7B,C). This is reminiscent of SMA disease progression in human patients who often develop muscle weakness in the legs prior to onset of in the arms.

Similar defects in leg function were observed upon examination of animals expressing Smn-RNAi #2 in neurons and muscles. Progressive loss of motor function was observed for all legs in aggregate (Fig 7D), and for individual leg pairs (Fig 7E,F). Additionally, we noted the same differential impact on individual leg pairs, with the anterior pair initially displaying much milder dysfunction than the most posterior pair (Fig 7E,F). None of these effects was observed for Oregon R controls reared and scored in the same manner (Fig 7D,E). Overall, the progressive loss of motor function and differential timing of affectation between posterior and anterior leg pairs suggests that these fly models of SMA are useful for modeling nuanced aspects of SMA onset and progression and suggest that highly specific mechanisms of SMA are conserved in fly.

## Conclusions and Discussion

We carried out a systematic assessment of disease-related phenotypes in a set of *Drosophila* SMA models at multiple developmental stages. This work was performed in larvae and adult flies expressing one of thirteen human patient-derived *Smn* missense mutations as well as flies expressing ubiquitous or neuromuscular-specific *Smn* knockdown. In larvae, we identified defects in developmental viability and locomotor function in the absence of overt muscle degeneration, leading to the conclusion that the larval stages are best suited to study of pre- and early onset stages of SMA. For the lines that reached the adult stage, we observed reduced longevity and locomotor defects with variable age of onset. Thus, the fly can be used to model progressive loss of motor function in SMA. We also report, for the first time in a model system, three alleles for late-onset (Type IV) SMA. Collectively, this set of SMA models provides a highly useful system for studying all stages of SMA pathogenesis across the full spectrum of disease severity.

### Modeling SMA in adult stage Drosophila

This study presents the first report of adult models of SMA in the fly, providing an opportunity to separate severe developmental complications from neuromuscular phenotypes. In typical cases, SMA patients appear healthy and normal at the time of birth with onset of impaired motor function occurring months or years after birth. Creating models that mimic this aspect of SMA has been difficult in the mouse, as most genetic manipulations that are sufficient to produce neuromuscular phenotypes also cause animals to be small, underdeveloped, and severely affected at the time of birth (Hsieh-Li *et al.*, 2000; Monani *et al.*, 2000). Pups for these models fail to grow significantly and die in the first weeks of life, reminiscent of Type 0 SMA in humans. Our Class III and IV fly models produce adults of normal size that appear otherwise healthy upon eclosion. These animals continue to mature as normal until they die between 1 and 7 weeks of life (control flies live for 12-13 weeks). Using animals that successfully progress through all developmental stages allows for separation of SMA phenotypes from gross developmental issues. This is important, as developmental arrest can independently affect mRNA processing and gene expression, nervous system development and growth, as well as many other phenotypes relevant to the study of SMA (Carrel *et al.*, 2006; McWhorter *et al.*, 2008; Garcia *et al.*, 2013; Garcia *et al.*, 2016; Perez-Garcia *et al.*, 2017). The ability to avoid these complications with genetic models that develop normally adds a valuable tool to the field of SMA research.

### Modeling intermediate forms of SMA

Mouse models for late-onset SMA have been difficult to generate. Copy number changes in human *SMN2* transgenes or C>T mutation of the endogenous mouse *Smn* allele cause dramatic shifts in phenotype from mild and largely unaffected to very severe with onset of symptoms *in utero* and death between 4-14 days after birth (Hsieh-Li *et al.*, 2000; Monani *et al.*, 2000). Addition of partially functional SMN-producing genes, such as *SMNΔ7* or *SMN2* leads to a similar bifurcation of phenotypes (Le *et al.*, 2005; Osborne *et al.*, 2012). Depending on the gene dose, mice are either strongly affected at birth or completely rescued throughout life. This binary relationship between SMN dosage and phenotype in mice has led to a dearth of robust mouse models of intermediate SMA. Consequently, the processes that precede onset of neuromuscular dysfunction and motor neuron death have been difficult to study in murine systems.

In the fly, we observe a more continuous range of phenotypic severity across many assays in the context of hypomorphic SMN levels and partial function. The mildest *Smn* missense mutations we assessed (D20V, G73R, and I93F) are phenotypically wild type at the time of eclosion. By 5 weeks of age, however, flies carrying these mutations begin to show locomotor dysfunction and expire after 1-3 additional weeks of life. Based on these observations, we assert that these three *Smn* missense mutations model moderate, later-onset Type III and IV SMA. These models will be a useful tool in examining the full range of SMA disease progression from pre-onset changes to organismal death in a single biological context.

### SMN missense mutations are partially functional in the absence of wild type SMN protein

Together with previous work (Praveen *et al.*, 2014; Garcia *et al.*, 2016; Gray *et al.*, 2018), the data presented here demonstrate that many SMN missense mutations retain partial function. As discussed above, the maternal contribution of SMN complicates this interpretation in early larval stages. By the start of the third instar larval stage, however, the maternal contribution is fully depleted and complete lack of SMN rapidly leads to developmental arrest and death. In our models, transgenic *Smn* is expressed from the native promotor throughout the life of the organism. In this context, we observe survival to differential degrees and developmental stages for each *Smn* missense mutations. These findings demonstrate that *Smn* missense mutations specifically modify the null phenotype and are inconsistent with the notion that they extend viability due to secondary effects caused by non-specific stabilization of the maternal contribution.

Furthermore, we successfully generated stable, viable fly stocks that exclusively express certain missense alleles of *Smn* in an otherwise *Smn* null background (*Flag-Smn^TG^, Smn^X7^/Smn^D^*). These animals are viable and fertile in the absence of any wild type SMN at any stage of their development, providing conclusive evidence that *Smn* missense mutations are at least partially functional. Given the conservation of SMN protein structure from invertebrates to vertebrates, we expect that this finding holds true in human patients as well. Our results contrast with those of a recent report using the SMNΔ7 mouse model, claiming that two mild patient-derived missense mutations (D44V and T274I) are non-functional in the absence of full-length SMN (Iyer *et al.*, 2018). This assertion is based on a single finding: that mice expressing only *SMN* missense mutations do not survive to birth. While this result clearly indicates that these two *SMN* missense mutations are not fully functional, it by no means rules out the possibility of partial function.

*Smn* null mice arrest at the blastula stage of embryonic development (Schrank *et al.*, 1997). Missense alleles that fully lack function would be expected to arrest at the same stage as the null allele. Unfortunately, neither this stage nor any other embryonic stage was assessed in the context of the D44V or T274I mutations to determine if *SMN* missense mutations partially rescue embryonic development (Iyer *et al.*, 2018). Notably, all of the human *SMN* missense alleles analyzed in the mouse to date have been expressed from randomly-integrated, variable copy number, cDNA-based transgenes (Gavrilina *et al.*, 2008; Workman *et al.*, 2009; Iyer *et al.*, 2018). Moreover, none of these studies uses a wild-type cDNA control, so it is difficult to compare the transgenic lines without the ability to control for expression levels, position effects, or secondary mutations caused by integration.

Indeed, transgenic rescue of embryonic lethality using human *SMN* constructs in the mouse is fraught with complication. Mice expressing two randomly-inserted cDNA copies of *SMN*Δ*7* die within 5 days of birth when crossed onto the FVB background (Monani *et al.*, 2000). However, this same *SMN*Δ*7* transgene completely fails to rescue embryonic lethality when crossed onto a C57/BL6 background (Gogliotti *et al.*, 2011; Osborne *et al.*, 2012; Meier *et al.*, 2018). Expressing one or two copies of human *SMN2* in an *Smn* null background is also insufficient to rescue embryonic lethality in mouse (Osborne *et al.*, 2012). These results are in contrast to the effects of manipulating SMN expression from the endogenous mouse locus. Mice bearing a C>T mutation in exon 7 of the endogenous *Smn* gene (mimicking human *SMN2*) are fully viable and fertile (Gladman *et al.*, 2010; Hammond *et al.*, 2010). Further, these mice live a normal lifespan when the C>T allele is present in either the homozygous or hemizygous state, displaying mild adult-onset phenotypes (Gladman *et al.*, 2010). This incongruity suggests that expression of randomly-inserted human *SMN* cDNAs is suboptimal in the context of restoring mouse viability. In fact, there is no evidence that a wild-type human *SMN* cDNA transgene can rescue mouse viability, as such a line has yet to be reported. For all of these reasons, additional work is needed in this area.

### Concordance between SMA severity and *Drosophila* phenotypes

The missense mutations examined here impact residues in the SMN protein that are well conserved from fly to human. Thus, we expect that mutating these residues will cause similar changes in SMN protein function and/or stability in both species. An indirect approach to assess this expectation is to compare the severity of SMA-related phenotypes in flies and SMA Type in humans. This analysis is somewhat complicated by several factors.

First, the diagnosis and classification of SMA has changed significantly over the 20 year period since the first *SMN1* missense mutations were reported. Thus, cross comparison of severity between patients diagnosed in different decades is imperfect. For example, a severe patient diagnosed with Type I SMA that was reported in the late 1990’s might be diagnosed as a Type 0 patient in the 2010’s. Similarly, a patient diagnosed as Type II two decades ago might be considered a Type III patient today.

Second, a major complicating factor is *SMN2* copy number. In several reports of missense mutations from the late 1990s, the *SMN2* copy number of the patient was neither assessed nor reported (Supp Table 2). Given the strong influence that *SMN2* copy number has on SMA severity, lack of this information prevents reasonable comparisons of human and fly SMA phenotypes. As mentioned above, genetic background is also known to modify SMA severity. Moreover, *SMN1* missense mutations are extremely rare and often reported in single patients or a pair of affected siblings whose disease severity is impacted by genetic background to an unknown degree.

From the perspective of the fly models, there are far fewer complicating genetic factors to consider. The fourteen Flag-tagged *Smn* lines described here were generated in a single genetic background, with a single copy of the transgene inserted at the identical chromosomal location. This approach allows for direct, reliable comparison of phenotypic severity between patient mutations in the fly. An important caveat is that fly development includes multiple body plans (larval vs adult) and stages of development (embryonic, larval, and pupal) prior to the adult stage and it is therefore difficult to directly compare to the relatively linear process of human development. For example, it is unclear how the onset of symptoms in larval stages compares to age of onset in human development. Due to this, phenotypic Class and SMA Type are not expected to align in terms of their number-designation. However, we do expect to see a correlation between human SMA Type and *Drosophila* Class designations, with the most severe human mutations also being the most severe in fly and vice versa.

Indeed, the information in Table 1 reveals that in almost every case, the severity observed in the fly is well aligned with human SMA Type. The one exception is the I93F/I116F mutant. Given that this mutation has only been observed in a single human patient, it is difficult to determine whether this residue may not be as functionally important for fly SMN as it is for human, or if confounding factors such as genetic background might enhance the SMA severity independent of *SMN1*.

**Table 1.**
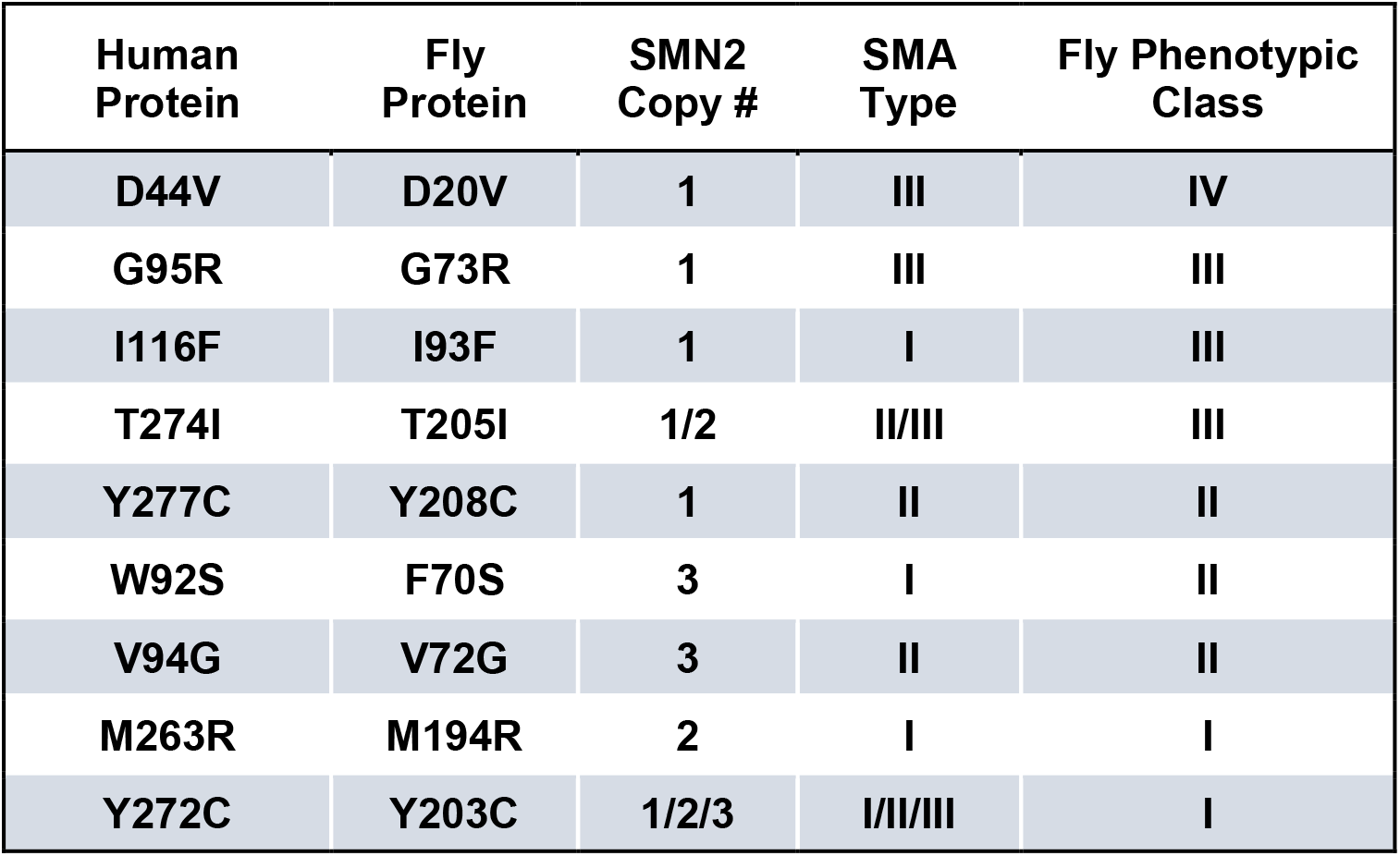
Alignment of phenotypic severity between human and fly. *Smn* missense mutations are shown in order of least severe to most severe, based on fly genotypes, with the corresponding information from human patients. The four missense mutations with unknown *SMN2* copy number (Y107C, G206S, G210C, and G210V, shown in Table 1) have been excluded here to prevent potentially inaccurate comparisons.

| Human Protein | Fly Protein | SMN2 Copy # | SMA Type | Fly Phenotypic Class |
| --- | --- | --- | --- | --- |
| D44V | D20V | 1 | III | IV |
| G95R | G73R | 1 | III | III |
| I116F | I93F | 1 | I | III |
| T274I | T205I | 1/2 | II/III | III |
| Y277C | Y208C | 1 | II | II |
| W92S | F70S | 3 | I | II |
| V94G | V72G | 3 | II | II |
| M263R | M194R | 2 | I | I |
| Y272C | Y203C | 1/2/3 | I/II/III | I |

In all other cases, we observe strong concordance between the phenotypic severity of fly and human. For example, the fly mutations Y203C and M194R cause the most severe phenotype in flies, phenocopying null alleles. The same is seen in human patients carrying Y272C and M263R mutations, which also phenocopy the *SMN1* null state in causing Type 1 SMA in the presence of 1 or 2 copies of *SMN2*. On the other end of the spectrum, the D44V, G95R, and T274I mutations appear to *improve* the SMA Type in patients, relative to the severity expected based on *SMN2* copy number. For example, humans hemizygous for both *SMN2* and *SMN1^T274^* display a relatively mild, Type II phenotype. Flies expressing the corresponding mutations (D20V, G73R, and T205I) are also mildly affected, with G73R and D20V causing adult-onset of locomotor defects. Overall, when appropriately considering SMA Type and *SMN2* copy number, we conclude that SMN missense mutations modeled in the fly faithfully recapitulate the human phenotypes.

---

**Supplemental Figure 1.**
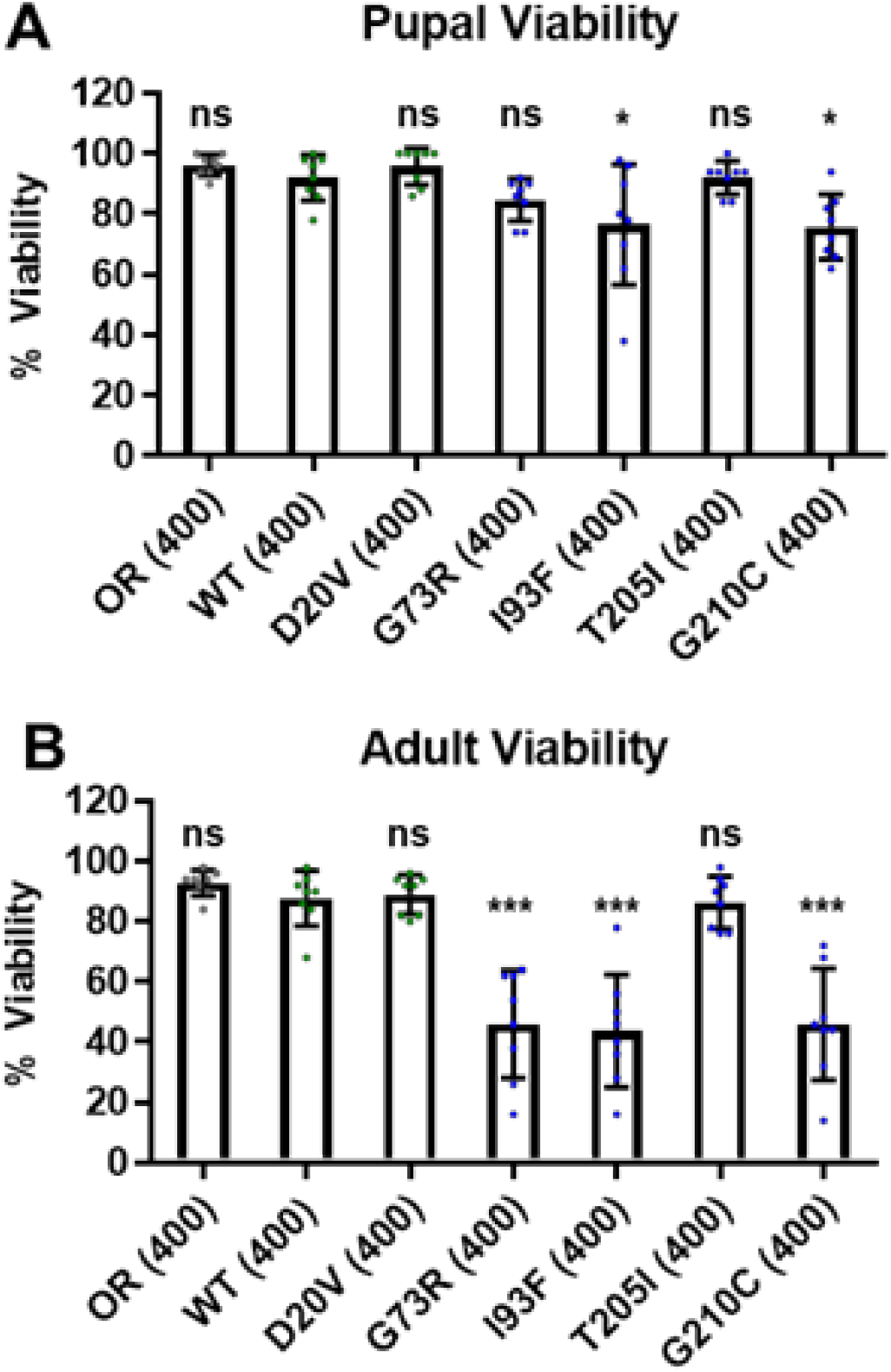
*Smn* missense mutations are sufficient for viability in the absence of maternally contributed wild type SMN. **A and B)** Developmental viability at the pupal stage (A) the adult stage (B) stable lines expressing only *Smn* missense mutations with maternal contribution of WT Smn present. **Data:** Bars show average. Error bars show standard error. **Data:** points represent biological replicates of 50 animals each, n-values (shown in parentheses next to genotypes) reflect the number of individual animals counted. **Statistical analysis:** Values above the data indicate significance vs WT from one-way ANOVA using the Dunnet correction for multiple comparisons. ns: not significant (p>0.05), * p<0.05 **p<0.01 ***p<0.001

**Supplemental Figure 2.**
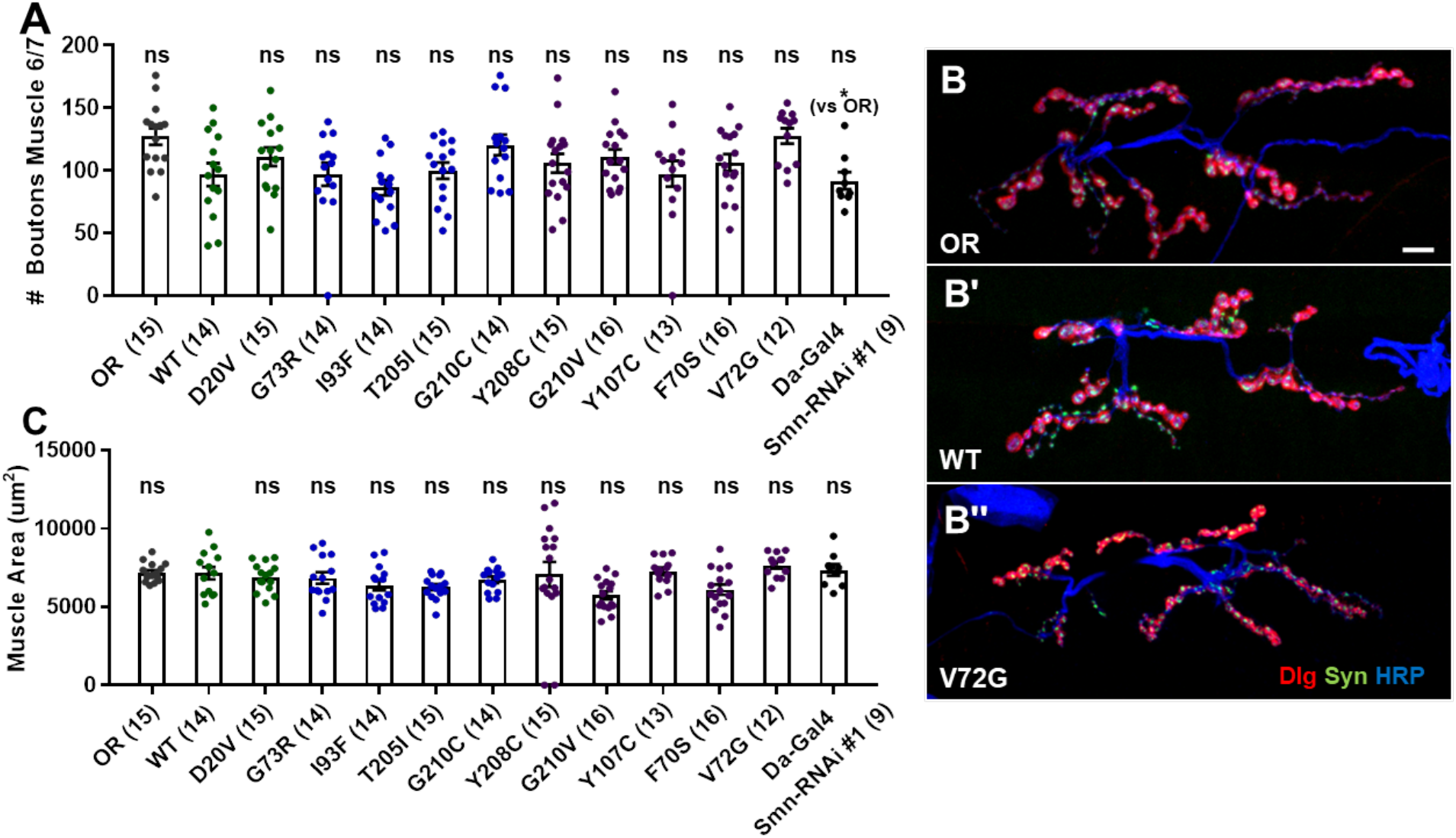
*Smn* missense mutations or knockdown has modest effects on neuromuscular junction (NMJ) structure. **A)** Bouton counts from the muscle 6/7 NMJ in wandering third instar larvae. **B)** Representative images for the data shown in A. Red marks Discs Large (Dlg), green marks synapsin (syn), and blue marks neuronal membranes. **C)** Muscle 6/7 combined area for the NMJs measured in A. **Data:** Bars show average. Error bars show standard error. Data points and n-values (shown in parentheses next to genotypes) reflect the number of individual neuromuscular junctions (A) or muscles (C) assayed. **Statistical analysis:** Values above the data indicate significance vs WT from one-way ANOVA using the Dunnet correction for multiple comparisons. ns: not significant (p>0.05), * p<0.05 **p<0.01 ***p<0.001

**Supplemental Figure 3.**
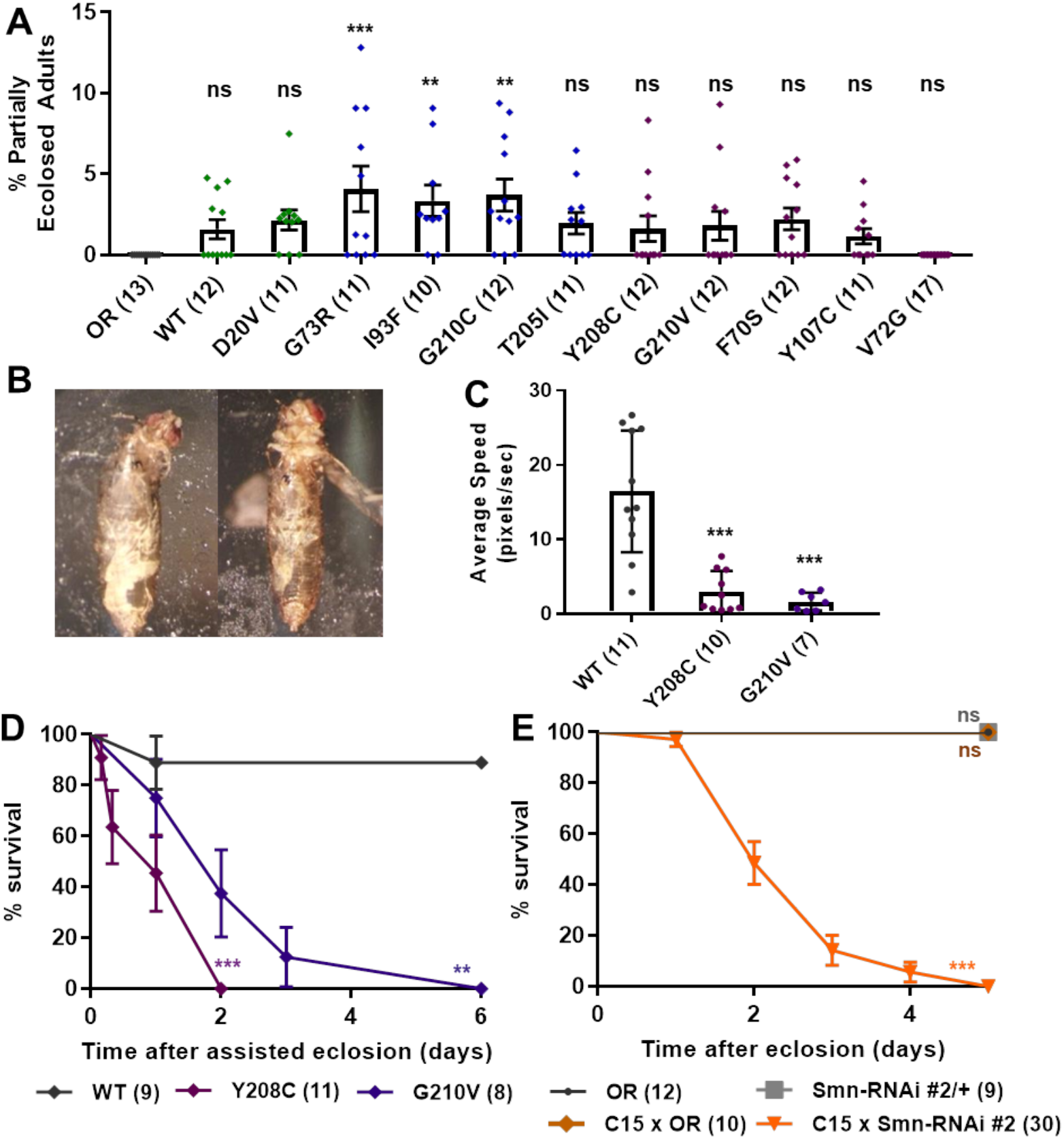
*Smn* missense mutants exhibit defects in eclosion. **A)** Percent of total larvae that complete pupal development but only partially eclose from their pupal case. **B)** Representative images of the partial eclosion phenotype quantified in A. **C)** Adult walking speed (pixels/second) for partially eclosed animals expressing the wild type *Smn* transgene or the Smn missense mutations Y208C or G210V. **D)** Survival curve for the same animals assayed in C. **E)** Survival curve for animals expressing neuromuscular *Smn* knockdown. **Data:** Bars show average. Error bars show standard error. Data points and n-values (shown in parentheses next to genotypes) reflect the number of individual animals assayed. **Statistical analysis:** For A and C, values above the data indicate significance vs WT from one-way ANOVA using the Dunnet correction for multiple comparisons. For D and E, values next to each survival curve represent p-values generated by Chi square analysis using the logrank rank/Mantel-Cox test. ns: not significant (p>0.05), * p<0.05 **p<0.01 ***p<0.001.

**Supplemental Table 1.** Information on all statistical analysis. (Table on next page) Statistical analysis was performed using GraphPad Prism 7 and includes corrections for multiple comparisons when appropriate.

| Figure Panel | Type of Data | Statistical Test | Correction for Multiple Comparison | F-value | p-value | R<sup>2</sup> |
| --- | --- | --- | --- | --- | --- | --- |
| 1A | Normal | One-way Anova | Dunnet | F(16,120) = 24.22 | 0.0001 | 0.7635 |
| 1B | Normal | One-way Anova | Dunnet | F(16,120) = 131.7 | 0.0001 | 0.9461 |
| 1C - Early | Normal | One-way Anova | Dunnet | F(13,156) = 90.39 | 0.0001 | 0.8828 |
| 1C - Late | Normal | One-way Anova | Dunnet | F(13,156) = 32.61 | 0.0001 | 0.7321 |
| 1C - Adult | Normal | One-way Anova | Dunnet | F(13,156) = 308.6 | 0.0001 | 0.9628 |
| Supp Fig 1A | Normal | One-way Anova | Dunnet | F(7, 56) = 8.78 | 0.0001 | 0.5232 |
| Supp Fig 2B | Normal | One-way Anova | Dunnet | F(7, 56) = 48.93 | 0.0001 | 0.8595 |
| 2A | Normal | One-way Anova | Dunnet | F(17, 671) = 54.38 | 0.0001 | 0.5794 |
| 2C | Normal | One-way Anova | Dunnet | F(17, 671) = 44.24 | 0.00001 | 0.5303 |
| 2D | Normal | One-way Anova | Dunnet | F(13, 545) = 21.68 | 0.0001 | 0.3426 |
| 2F | Normal | One-way Anova | Dunnet | F(13, 545) = 3.545 | 0.0001 | 0.0780 |
| Supp Fig 2A | Normal | One-way Anova | Dunnet | F(14,195) = 2.369 | 0.0046 | 0.1454 |
| Supp Fig 2C | Normal | One-way Anova | Dunnet | F(14, 194) = 1.583 | 0.0865 | 0.1026 |
| 3A | Normal | One-way Anova | Dunnet | F(8, 55) = 16.31 | 0.0001 | 0.7035 |
| 3B | Normal | One-way Anova | Dunnet | F(8, 55) = 64.07 | 0.0001 | 0.9031 |
| 3C | Normal | One-way Anova | Dunnet | F(8, 179) = 14.03 | 0.0001 | 0.3853 |
| 4A | Survival data | Logrank/Mantel-Cox Test | Bonferroni | Corrected significance at p<0.0026 |  |  |
| 4B | Survival data | Logrank/Mantel-Cox Test | Bonferroni | Corrected significance at p<0.0026 |  |  |
| 4C | Survival data | Logrank/Mantel-Cox Test | Bonferroni | Corrected significance at p<0.0026 |  |  |
| 4F | Survival data | Logrank/Mantel-Cox Test | Bonferroni | Corrected significance at p<0.0056 |  |  |
| 4G | Survival data | Logrank/Mantel-Cox Test | Bonferroni | Corrected significance at p<0.0056 |  |  |
| 5A | Normal | One-way Anova | Dunnet | F(11, 1242)= 21.36 | 0.0001 | 0.1591 |
| 5C | Normal | One-way Anova | Dunnet | F(5, 157) = 9.473 | 0.0001 | 0.2318 |
| 5D | Normal | One-way Anova | Dunnet | F(5, 404) = 52.56 | 0.0001 | 39.4100 |
| 6B | Normal | One-way Anova | Dunnet | F(3, 256) = 6.607 | 0.0003 | 0.0719 |
| 6C | Normal | One-way Anova | Dunnet | F(3, 192) = 13.39 | 0.0001 | 0.1730 |
| Supp Fig 3A | Normal | One-way Anova | Dunnet | F(13, 157)= 4.06 | 0.0001 | 0.2514 |
| Supp Fig 3C | Normal | One-way Anova | Dunnet | F(2, 25) = 22.01 | 0.0001 | 0.6378 |
| Supp Fig 3D | Survival data | Logrank/Mantel-Cox Test | Bonferroni | Corrected significance at p<0.025 |  |  |
| Supp Fig 3E | Survival data | Logrank/Mantel-Cox Test | Bonferroni | Corrected significance at p<0.012 |  |  |
| 7A - 4 hrs | Normal | One-way Anova | Dunnet | F(2, 70) = 29.71 | 0.0001 | 0.4591 |
| 7A - 8 hrs | Normal | One-way Anova | Dunnet | F(2, 67) = 45.55 | 0.0001 | 0.5762 |
| 7A -12 hrs | Normal | One-way Anova | Dunnet | F(2, 73) = 62.00 | 0.0001 | 0.6294 |
| 7A - 1 day | Normal | One-way Anova | Dunnet | F(2, 73) = 97.65 | 0.0001 | 0.7279 |
| 7A - 2 days | Normal | One-way Anova | Dunnet | F(2, 73) = 129 | 0.0001 | 0.7795 |
| 7A - 3 days | Normal | One-way Anova | Dunnet | F(2, 79) = 156.6 | 0.0001 | 0.7988 |
| 7A - 4 days | Normal | Student's T-test | N/A | F(23, 24) = 1.98 | 0.0001 | 0.7659 |
| 7A -5 days | Normal | Student's T-test | N/A | F(23, 24) = 1.96 | 0.0001 | 0.7679 |
| 7A - 6 days | Normal | Student's T-test | N/A | F(23, 24) = 1.99 | 0.0001 | 0.9233 |
| 7E - 8 hrs | Normal | Student's T-test | N/A | F(89, 35) = 1.99 | 0.0001 | 0.3959 |
| 7E - 24 hrs | Normal | Student's T-test | N/A | F(89, 35) = 1.99 | 0.0001 | 0.2085 |
| 7E - 48 hrs | Normal | Student's T-test | N/A | F(89, 35) = 1.99 | 0.0001 | 0.5020 |
| 7E - 72 hrs | Normal | Student's T-test | N/A | F(89, 35) = 1.99 | 0.0001 | 0.6336 |
| 7E - 96 hrs | Normal | Student's T-test | N/A | F(89, 35) = 1.99 | 0.0001 | 0.6336 |

**Supplemental Table 2.** SMN missense mutation information from human patients and fly. When multiple SMA types and *SMN2* copy numbers are present, the order of the information for each criterion corresponds such that the first SMA type shown corresponds to the first *SMN2* copy number listed for a given mutation and so on and forth.

| Human Mutation | Human Protein | Fly Protein | Protein Domain | # Patients Reported | Human SMA Type | SMN2 Copy # | Fly Phenotypic Class | Primary Reference |
| --- | --- | --- | --- | --- | --- | --- | --- | --- |
| c.131A>T | D44V | D20V | Gemin2 binding | 1 | III | 1 | IV | Sun et al., 2005 |
| c.275G>C | W92S | F70S | Tudor domain | 2 | I | 3 | II | Kotani et al., 2007 |
| c.281T>G | V94G | V72G | Tudor domain | 1 | II | 3 | II | Clermont et al., 2004 |
| c.283G>C | G95R | G73R | Tudor domain | 1 | III | 1 | III | Sun et al., 2005 |
| c.346A>T | I116F | I93F | Tudor domain | 1 | I | 1 | III | Cusco et al., 2004 |
| c.389A>G | Y130C | Y107C | Tudor domain | 1 | NR | NR | II | Prior, 2007 |
| c.788T>G | M263R | M194R | YG box | 1 | I | 2 | I | Clermont et al., 2004 |
| c.815A>G | Y272C | Y203C | YG box | 11 | I/II/III | 1/2/3 | I | Wirth et al., 1999 |
| c.821C>T | T274I | T205I | YG box | 4 | II/III | 1/2 | III | Wirth et al., 1999 |
| c.856G>C | G275S | G206S | YG box | 1 | III | NR | I | Skordis et al 2001 |
| c.830A>G | Y277C | Y208C | YG box | 1 | II | 1 | II | Yamamoto et al. 2013 |
| c.868G>T | G279C | G210C | YG box | 2 | II/III | NR | III | Wang et al., 1998 |
| c.869G>T | G279V | G210V | YG box | 2 | I | NR | II | Talbot et al., 1997 |

## Acknowledgements

This work was supported by a grant from the USA National Institute of General Medical Sciences (NIGMS), R01-GM118636 (to A.G.M.). A.M.S. was supported by a Seeding Postdoctoral Innovators in Research and Education (SPIRE) fellowship from the National Institutes of Health (NIH), K12-GM000678 (to D.T. Lysle). We thank C.A. Frank for providing the C15 Gal4 driver line. Note, a version of this manuscript has been posted on a pre-print server (Spring et al. 2018, doi: https://doi.org/10.1101/394908).

